# Microsaccadic suppression of peripheral perceptual detection performance as a function of foveated visual image appearance

**DOI:** 10.1101/2024.03.07.583880

**Authors:** Julia Greilich, Ziad M. Hafed

## Abstract

Microsaccades are known to be associated with a deficit in perceptual detection performance for brief probe flashes presented in their temporal vicinity. However, it is still not clear how such a deficit might depend on the visual environment across which microsaccades are generated. Here, and motivated by studies demonstrating an interaction between visual background image appearance and perceptual suppression strength associated with large saccades, we probed peripheral perceptual detection performance of human subjects while they generated microsaccades over three different visual backgrounds. Subjects fixated near the center of a low spatial frequency grating, a high spatial frequency grating, or a small white fixation spot over an otherwise gray background. When a computer process detected a microsaccade, it presented a brief peripheral probe flash at one of four locations (over a uniform gray background), and at different times. After collecting full psychometric curves, we found that both perceptual detection thresholds and slopes of psychometric curves were impaired for peripheral flashes in the immediate temporal vicinity of microsaccades, and they recovered with later flash times. Importantly, the threshold elevations, but not the psychometric slope reductions, were stronger for the white fixation spot than for either of the two gratings. Thus, like with larger saccades, microsaccadic suppression strength can show a certain degree of image-dependence. However, unlike with larger saccades, stronger microsaccadic suppression did not occur with low spatial frequency textures. This observation might reflect the different spatio-temporal retinal transients associated with the small microsaccades in our study versus larger saccades.

## Introduction

Microsaccades are small saccades that occur periodically during attempted gaze fixation (Rolfs, 2009). These eye movements are similar to larger scanning saccades in both kinematics and underlying neurophysiological mechanisms (Hafed, 2011; Hafed, Goffart, & Krauzlis, 2009; Hafed & Krauzlis, 2012; Zuber, Stark, & Cook, 1965). Moreover, microsaccades aid in optimizing gaze position during foveal visually-guided behavior (J. Bellet, Chen, & Hafed, 2017; Ko, Poletti, & Rucci, 2010; Poletti, Listorti, & Rucci, 2013; Shelchkova, Tang, & Poletti, 2019; Tian, Yoshida, & Hafed, 2016, 2018), just like larger saccades allow targeting new scene locations for detailed visual analysis. Microsaccades may thus be thought of as scanning eye movements, but on the small scale of foveal vision (Hafed, Chen, Tian, Baumann, & Zhang, 2021).

Because of the similarity between microsaccades and saccades, it is expected that both types of eye movements may similarly affect visual processing. Indeed, some of the earliest studies on the well-known phenomenon of perceptual saccadic suppression were first conducted with microsaccades (Beeler, 1967; Zuber & Stark, 1966). In this phenomenon, the detection of brief peri-saccadic or peri-microsaccadic visual stimulus onsets is strongly impaired. Later work characterized the neural correlates of this phenomenon, again with microsaccades. Specifically, and using a similar approach to that used in psychophysical studies (that is, by presenting visual onsets around the time of microsaccade onset), it was found that visual neural sensitivity in multiple brain areas can be suppressed for peri-microsaccadic stimulus events (Chen & Hafed, 2017; Chen, Ignashchenkova, Thier, & Hafed, 2015; Hafed & Krauzlis, 2010). These studies complemented other studies demonstrating an influence of both saccades and microsaccades on ongoing visual neural activity in the absence of sudden stimulus onsets (Bosman, Womelsdorf, Desimone, & Fries, 2009; Crowder, Price, Mustari, & Ibbotson, 2009; Herrington et al., 2009; Kagan, Gur, & Snodderly, 2008; Leopold & Logothetis, 1998; Reppas, Usrey, & Reid, 2002; Sylvester, Haynes, & Rees, 2005; Thiele, Henning, Kubischik, & Hoffmann, 2002; Wurtz, 1968, 1969a, 1969b).

With large saccades, the phenomenon of perceptual saccadic suppression has been extensively studied for several decades. While the underlying mechanisms for this phenomenon may not be fully elucidated yet, increasing evidence suggests that saccadic suppression arises through a potential interaction between the movement command for saccade generation itself and the viewed visual image properties (Baumann, Idrees, Munch, & Hafed, 2021; Braun, Schutz, & Gegenfurtner, 2017; Bremmer, Kubischik, Hoffmann, & Krekelberg, 2009; Brooks, Impelman, & Lum, 1981; Duffy & Lombroso, 1968; Gremmler & Lappe, 2017; Idrees, Baumann, Franke, Munch, & Hafed, 2020; Mackay, 1970; Maij, Matziridi, Smeets, & Brenner, 2012; Matin, Clymer, & Matin, 1972; Mitrani, Mateeff, & Yakimoff, 1971; Mitrani, Yakimoff, & Mateeff, 1973; Riggs & Manning, 1982; Wurtz, 2008). In fact, the dependence of perceptual saccadic suppression on the properties of the viewed visual image already starts in the retina, the very first visual processing stage after the eye optics (Idrees et al., 2020; Idrees et al., 2022). Interestingly, if motor commands of the superior colliculus do contribute to saccadic suppression via ascending corollary discharge projections (Berman, Cavanaugh, McAlonan, & Wurtz, 2017; Berman & Wurtz, 2011; Isa & Hall, 2009; Lee et al., 2007; Phongphanphanee et al., 2011), then an interaction between saccade movement commands and underlying visual image properties for saccadic suppression becomes almost inevitable, especially given the discovery of visual sensory tuning within the superior colliculus motor bursts themselves (Baumann, Bogadhi, Denninger, & Hafed, 2023; Zhang, Malevich, Baumann, & Hafed, 2022).

Having said that, most prior work on dependencies of perceptual saccadic suppression on visual image appearance has focused on large saccades. Thus, here, we aimed to document potential changes in the properties of microsaccadic suppression of perceptual detection performance as a function of what image was being fixated when a microsaccade was generated. Using an approach similar to that we used recently (Idrees et al., 2020), we measured peripheral perceptual detection thresholds and sensitivity (slopes of psychometric curves) when microsaccades were generated over a stable background image. We found that both detection thresholds and sensitivity were generally impaired perimicrosaccadically, but that only threshold elevations (and not sensitivity reductions) showed a dependence on the visual appearance of the image that was foveated. Moreover, the visual dependence of threshold elevations was decidedly different from that expected from our earlier results with larger saccades (Idrees et al., 2020), possibly reflecting the largely different spatio-temporal scales with which microsaccades and saccades modulate retinal image statistics (Mostofi et al., 2020).

## Methods

### Subjects and ethical approvals

We recruited 12 human subjects for this study. Of these, seven were female, and five were male. The subjects were aged 21-42 years, and each took part in three experimental sessions. The first and last sessions included 690 trials each, and the second session included 675 trials. Two subjects had too few baseline trials (defined explicitly in more detail below), so they each performed an additional session of 600 trials. The subjects individually took up to three short breaks during each session. All subjects consented to the experiment, and the procedures were approved by ethical committees of the Medical Faculty of the University of Tübingen. The subjects, who were naive to the purposes of the study, were also compensated financially for their time.

### Laboratory setup

All experiments were performed in the same setup as that used for recent studies (Baumann et al., 2021; Idrees et al., 2020; Idrees et al., 2022). In brief, the subjects sat comfortably 57 cm in front of a CRT display spanning approximately +/- 17 deg horizontally and +/- 13 deg vertically. The display had a refresh rate of 85 Hz and a pixel resolution of 41 pixels/deg. The display was also calibrated and linearized for luminance, as we used a similar procedure of collecting psychometric curves as in our earlier studies on perceptual thresholds (Baumann et al., 2021; Idrees et al., 2020). The room was otherwise dark.

We tracked eye movements using a video-based eye tracker (EyeLink 1000, SR Research), which was desk-mounted. This required stabilizing the head position, which we did using a custom-built device involving a chin rest, a forehead rest, constraints around the temple of the head, and a head band wrapped behind the head (Hafed, 2013).

We controlled the experiments using the Psychophysics Toolbox (Brainard, 1997; Kleiner, Brainard, & Pelli, 2007; Pelli, 1997) and Eyelink Toolbox (Cornelissen, Peters, & Palmer, 2002).

### Stimuli

We asked the subjects to always maintain fixation on an image that was presented at the center of the display in every trial. Across trials, three different images were possible. The first two were vertical gabor gratings, and the last was a small, white fixation spot. The white fixation spot was a square of 0.12 deg and 94.91 cd/m^2^ luminance. The gabor gratings could have a spatial frequency of either 0.5 cycles/deg (referred to as the low spatial frequency grating) or 5 cycles/deg (referred to as the high spatial frequency grating). These gabors had a σ parameter of 1.75 deg, and they thus visually spanned approximately +/- 6 deg horizontally and vertically on the display. The underlying sine wave luminance of each grating had a contrast of 100%, and we randomly picked one phase on every trial (from 8 possible phases equally spaced between 0 and 2π). The gray background luminance was 22.15cd/m^2^.

To probe perceptual thresholds, we also presented brief flash stimuli, which were squares of 1 deg size (each trial was associated with only one single probe flash presentation, as described in more detail below). These probe stimuli were presented at 9.1 deg from the screen center either horizontally or vertically, and their luminance varied across trials. This allowed us to collect full psychometric curves of detection performance. Specifically, each flash probe had a luminance increment above the background screen luminance of 2, 4, 5, 6, 8, 10, 11, or 14 computer register steps, with each step causing an actual luminance increase of 0.56 cd/m^2^ (which we measured after display calibration). The range of luminance steps used in any given condition depended on the time at which we presented the probe flash relative to a detected microsaccade (see details below). Specifically, our experiments involved gaze-contingent microsaccade detection (Baumann et al., 2021; Chen & Hafed, 2013; Idrees et al., 2020), and we expected higher perceptual thresholds for flashes very close to microsaccade onset than for later flashes. Thus, we used luminance increment steps of 2, 5, 8, 11, and 14 for the trials with higher expected perceptual thresholds, and we used 2, 4, 6, 8, and 10 increment steps for the trials with flashes farther away from microsaccades. In post-hoc analyses (see below), we redetected microsaccades and reclassified flash times, and we used all available luminance increments of a given trial type for fitting the psychometric curves.

### Experimental procedures

Each session took approximately 50-60 minutes of time. In each trial, a central image first appeared (low spatial frequency grating, high spatial frequency grating, or white fixation spot). The subjects were asked to maintain fixation on the image. For the white fixation spot, this was easy because it was the only visible item on the display, and it was very small. For the gratings, we instructed the subjects to look somewhere near the middle of the image. This allowed equalizing the eccentricity of the probe flashes across their four possible locations (also see Results). After 250 ms from image onset, we started a computer process of monitoring eye positions in real-time (Baumann et al., 2021; Idrees et al., 2020). If a microsaccade was detected between 250 ms and 700 ms from the image onset, we presented a probe flash for only one display frame at one of four possible locations (as described above). Moreover, the probe flash was triggered at 0, 25, or 75 ms after online eye movement detection. If no microsaccade was detected at all by 700 ms, we presented the probe flash anyway at one of the four locations. Later, in post-hoc analyses, we checked for the time of the nearest microsaccade to flash onset in this case, and we classified the trial according to our standard time course analyses (see more data analysis details below). After the probe flash, the computer waited for the subject to press one of four buttons, indicating the perceived flash location (right, left, up, or down). If no button was pressed after 1 second, a text message appeared on the display asking the subject to respond (and guess the flash location if necessary).

Our process for online microsaccadic eye movement detection was described earlier (Chen & Hafed, 2013) and successfully used for both microsaccades (Chen & Hafed, 2013) and larger saccades (Baumann et al., 2021; Idrees et al., 2020). Briefly, in every millisecond, we collected a running series of the latest 5 ms of eye position samples. Within each collection, we estimated the rate of change of eye position by fitting a line to the collected samples. To reduce effects of noise, we then took the median of the latest 3 slope measurements and flagged a microsaccade occurrence if the value of the slope was larger than a user-adjusted threshold. Note that this procedure necessarily delayed our estimate of microsaccade onset. This is why we redetected all microsaccades in later offline analyses after data collection, and we then recalculated probe flash times to actual microsaccade onset times. Also note that with this approach, we did not systematically measure pre-microsaccadic perceptual performance. This was the case because it would have required excessively more trials: without experimental control on microsaccade onset time, collecting pre-microsaccadic performance would entail presenting flashes at random times and then collecting enough trials to catch ones in which the flashes occurred pre-microsaccadically; with full psychometric curves (requiring repeated presentations of a given flash luminance), this requires much more data sessions per subject. In our previous work, we reached qualitatively similar conclusions whether we used our current approach or one also including pre-saccadic perceptual suppression trials (Idrees et al., 2020).

### Data analysis

We only analyzed trials with button press reaction times between 300 and 3000 ms. We also only included trials in which there were no flagged eye position samples within +/- 250 ms from probe flash onset. Flagged eye position samples could occur due to blinks (missing eye position data) or eye tracker noise (for example, by interference from eye lashes if subjects started squinting).

We detected all microsaccades using our established methods (M. E. Bellet, Bellet, Nienborg, Hafed, & Berens, 2019; Chen & Hafed, 2013). We then recalculated probe flash times relative to the recalculated microsaccade onset times. This accounted for the fact that online microsaccade detection was necessarily always slightly later than actual microsaccade onset (due to the data buffering mentioned above). Trials with saccades near probe flash onset that were larger than 3 deg in radial amplitude were excluded. These were extremely rare; in fact, for all three image types, most microsaccades were less than 1 deg in amplitude (see Results).

To obtain a time course of microsaccade-related perceptual threshold elevations (immediately around microsaccades) followed by recovery (for longer latency probe flash times relative to saccade onset), we then classified all trials into three different groups according to the time of the closest microsaccade to flash onset. The first group included all trials in which the closest microsaccade to flash onset started within +/- 50 ms from the probe flash event (because of our reclassification of microsaccade onset times in post-hoc analyses, there could be very few trials with a flash right before a microsaccade, and that is why we included 50 ms on either side of flash onset time here). This group of trials was expected to be associated with impaired detection performance (and it was called the group containing the microsaccadic suppression time bin). The second group included all trials in which the flash occurred 70 to 150 ms after the onset of the closest microsaccade to the flash. This group of trials was expected to show recovery in perceptual thresholds (and the time bin associated with it was called the recovery time bin). Finally, the third group of trials was that in which there were no microsaccades at all within +/- 250 ms from probe flash onset. These trials were called the baseline trials.

We further filtered trials according to microsaccade direction. Specifically, we found that the great majority of microsaccades were predominantly horizontal (see Results), consistent with earlier observations (Engbert & Kliegl, 2003; Laubrock, Engbert, & Kliegl, 2005). Therefore, we only included trials in the analyses for which the closest microsaccade to probe flash onset (including the three temporal categorizations described above) was predominantly horizontal (directional angle within <45 deg from the horizontal direction). This was reasonable given the vertical gratings, which would mean that horizontal microsaccades would be expected to cause the largest sensory transients in the brain after they occur (Khademi, Chen, & Hafed, 2020).

To assess perceptual performance, for each subject and time bin, we plotted the proportion of correct trials as a function of probe flash luminance increment above the background luminance. We then fit psychometric curves using the *psignifit 4 toolbox* (Schutt, Harmeling, Macke, & Wichmann, 2016); we specifically used the cumulative gaussian function for fitting. We defined the perceptual threshold as the luminance increment of the probe flash yielding 62.5% correct performance rate (given that ours was a 4-alternative forced choice paradigm with a 25% chance performance rate). For each subject and time bin, we estimated the threshold, and we then compared thresholds between conditions (e.g. low spatial frequency versus white fixation spot in the suppression time bin; or the suppression time bin versus the baseline time bin) across the population by showing mean and SEM across all subjects.

We performed statistical tests using the Friedman non-parametric test. Specifically, to test for an impact of probe flash time within a given image condition, we compared thresholds across subjects by grouping the measurements into three time bins as the factors of the statistical analysis (suppression, recovery, and baseline time bins). If the Friedman test had a p-value of less than 0.05, we then performed pairwise Wilcoxon signed rank tests to check which factors (time bins) were associated with thresholds that were different from each other. We consider an alpha value of less than 0.05 as significant in this study. For checking whether the threshold depended on the foveated image type, we again performed a Friedman test, but only on data from within a given time bin (e.g. the microsaccadic suppression time bin). This time, the factors of the statistical test were the three image types. We then used the same logic of post-hoc pairwise comparisons. We report all p-values in Results.

We also assessed sensitivity at threshold by measuring the slope of the psychometric curve near the probe flash levels causing threshold performance. To do so, for each curve, we divided the difference in performance (65% minus 60% correctness rate) by the difference in luminance values yielding 65% and 60% correctness rate. This gave an estimate of the slope of the psychometric curve at threshold. We then performed similar statistical tests on the slope measurements as we did for the detection thresholds.

For comparing microsaccade amplitudes as a function of foveated image type, we first averaged the microsaccade amplitude per condition within each subject’s data. Then, we visualized the population results by averaging across subjects, and showing SEM ranges. We also did this for microsaccade peak velocities, in order to assess the movements’ kinematics.

Finally, we also characterized where subjects fixated their gaze at the time of probe flash presentation. To do so, we averaged eye position in the interval from −25 ms to +75 ms relative to probe flash time. Then, we plotted the measurements across trials for each fixated image time. We picked more time samples after the flash than before because the great majority of flashes were triggered (by design of the experiment) after a microsaccadic event, and we wanted to avoid including the microsaccadic displacement itself in the eye position measurement.

## Results

We asked subjects to fix their gaze near the center of one of three possible images (Fig. 1A-C). One image was a low spatial frequency vertical gabor grating of 0.5 cycles/deg spatial frequency (Fig. 1A); the other was a high spatial frequency vertical gabor grating of 5 cycles/deg spatial frequency (Fig. 1B); and the third was a small white fixation spot (Fig. 1C). We then presented a brief probe flash peripherally for just one display frame (at one of the four cardinal directions; 9.1 deg eccentricity), and we asked subjects to indicate where it appeared on the display (4-alternative forced choice paradigm) (Fig. 1A-C). The probe flash was designed to appear at different time intervals relative to the occurrence of a microsaccade (Methods; examples shown in Fig. 1D-F), and the microsaccade itself was not explicitly instructed. Rather, the subjects were only told to look at the center of the image, and the computer waited for online microsaccade detection in order to trigger the probe flash (Methods).

**Figure 1.**
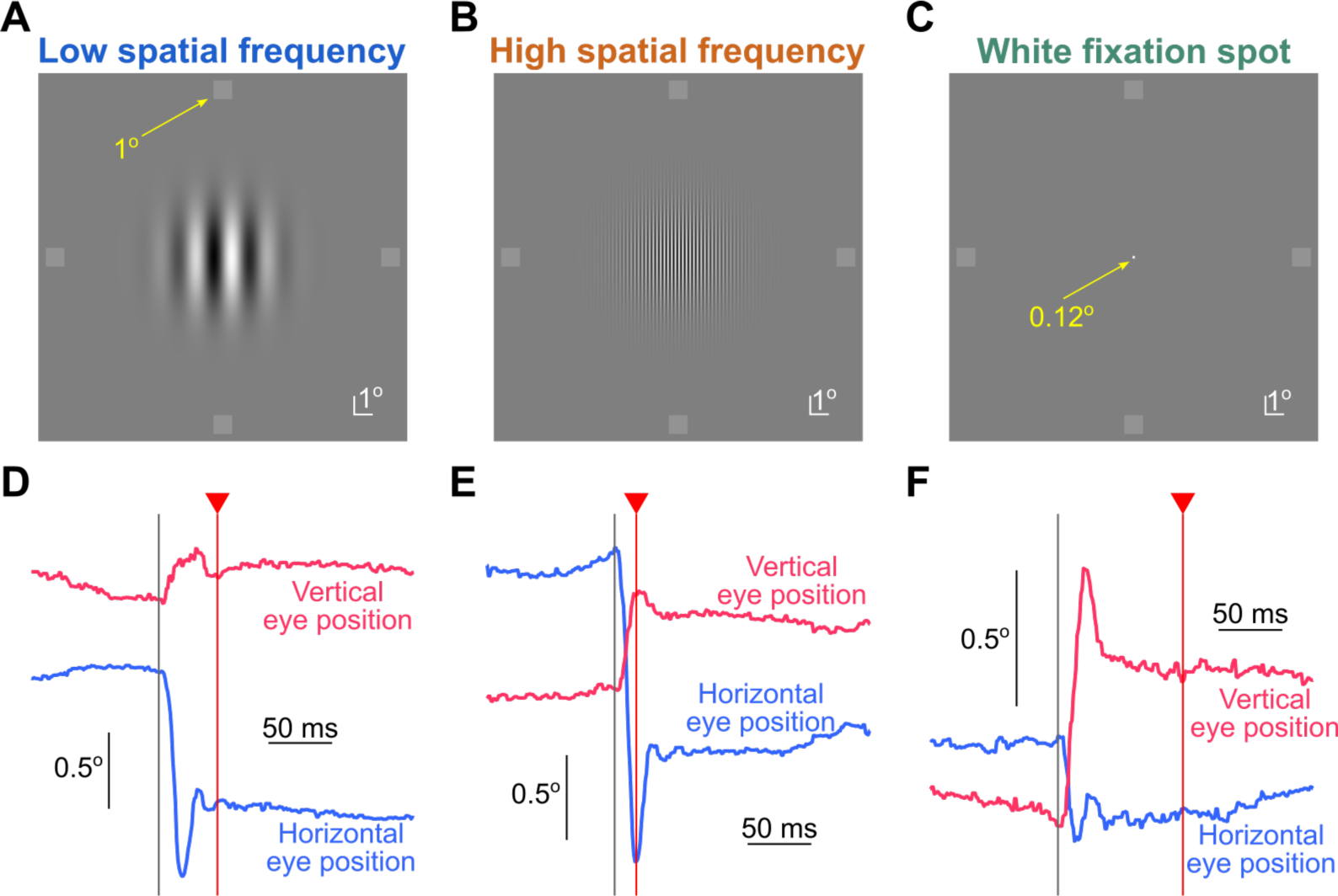
Experimental paradigm. (A-C) The images that were fixated in this study (Methods), along with the possible brief probe flash locations (small dim squares in the periphery). In every trial, the subjects were instructed to fixate near the center of the image. At a variable time from online microsaccade detection, a brief probe flash was presented at one of four peripheral locations (right, left, up, or down from the image center; we show the four probes here simultaneously only for illustration purposes because only one flash was presented per trial). The subjects had to indicate where they saw the brief probe flash. The luminance of the probe flash varied from trial to trial, in order to collect full psychometric curves. **(D-F)** Example eye position traces and probe flash onset times (vertical red lines) from one example subject (S05) in each of the image conditions shown in **A**-**C**. An upward deflection in the shown eye position traces indicates a rightward gaze shift for horizontal eye position and an upward gaze shift for vertical eye position. Our goal was to assess perceptual detection performance as a function of the time of flash onset relative to microsaccade onset, and also as a function of the different underlying foveated images. Note how the probe flash time was variable relative to microsaccade onset across different trials.

Across all trials, in post-hoc analyses, we searched for the nearest microsaccade to probe flash onset. We then first assessed the metric and kinematic properties of these eye movements. Independent of the underlying foveated image, the majority of microsaccades that occurred were predominantly horizontal (Fig. 2). For the grating images, this likely reflected the vertical orientation of the gratings, since orthogonal eye movements to the luminance gradient would be expected to give rise to the most useful information to the visual system about the underlying image (Rucci, Iovin, Poletti, & Santini, 2007). This is also consistent with neurophysiological signatures of microsaccade-induced visual reafferent responses at extra-foveal eccentricities, in which orthogonal eye movements scaled to a given spatial frequency give rise to the clearest modulations (Hafed, Chen, & Khademi, 2022; Khademi et al., 2020). For the white fixation spot, there were slightly more vertical eye movements than with the gabor gratings (likely reflecting the square appearance of the fixation spot, which includes both horizontal and vertical edges); nonetheless, the overall predominantly horizontal signature of eye movement directions with the white fixation spot was consistent with previous reports (Engbert & Kliegl, 2003; Laubrock et al., 2005). All of these observations led us to focus our remaining analyses on predominantly horizontal eye movements with an absolute direction from horizontal of less than 45 deg; we obtained generally similar results when we included all trials into the analyses, as expected given the large number of predominantly horizontal movements seen in Fig. 2. It is also interesting to note here that there were barely any downward microsaccades in our data at all (Fig. 2); this might be related to general tendencies of the oculomotor system to bias gaze upward, whether in visual or memory conditions (Goffart, Hafed, & Krauzlis, 2012; Goffart, Quinet, Chavane, & Masson, 2006; Khademi et al., 2024; Malevich, Buonocore, & Hafed, 2020; Snodderly, 1987; White, Sparks, & Stanford, 1994; Willeke, Cardenas, Bellet, & Hafed, 2022; Zelinsky, 1996).

**Figure 2.**
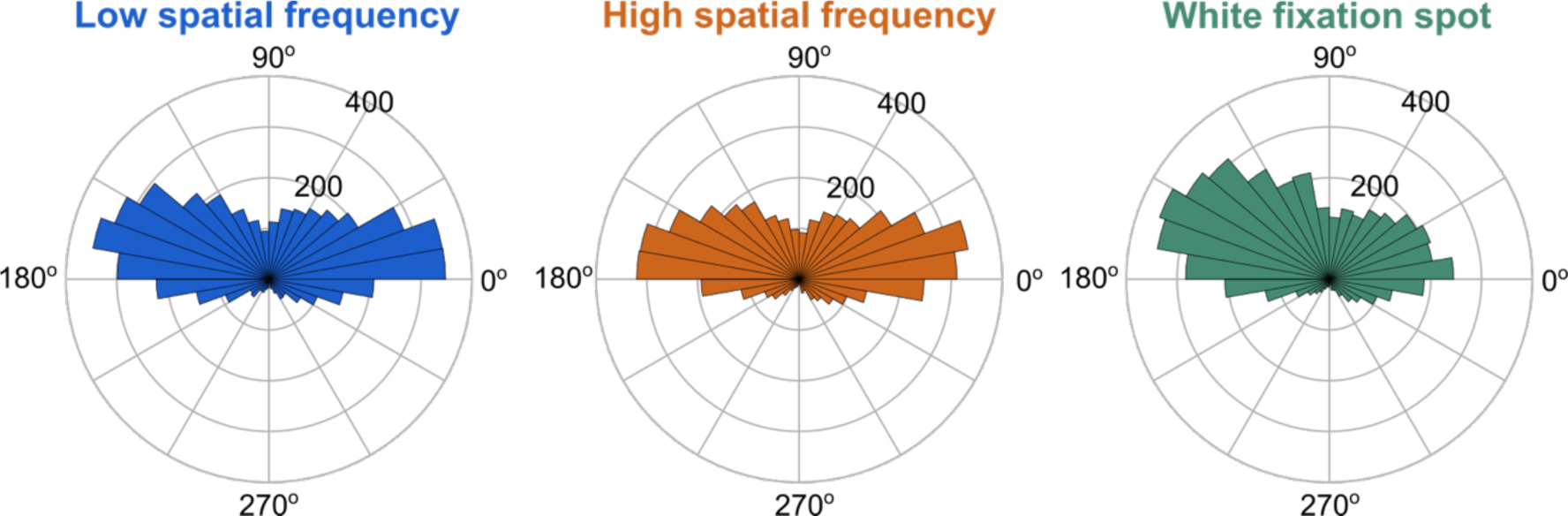
Predominantly horizontal nature of microsaccades in our experiments. Each panel shows the direction distribution of observed microsaccades (closest to probe flash onset time in every trial; Methods) for each fixated image of Fig. 1 (pooled across all subjects). As can be seen, most movements were predominantly horizontal. There was also a significant paucity of downward microsaccades in all conditions.

In terms of movement amplitudes, we found that predominantly horizontal microsaccades tended to be very slightly larger for the low spatial frequency image than for the high spatial frequency image, and microsaccades were also the smallest in size for the small white fixation spot. These results can be seen in Fig. 3A, and they are consistent with the above-mentioned ideas about how fixational eye movement properties can be strategically optimized by the visual-oculomotor system to maximize information gain from the underlying images. Nonetheless, in all cases, the microsaccades that we investigated in this study were always significantly smaller than 1 deg in radial amplitude regardless of the underlying image type (Fig. 3A), and they also obeyed the main sequence relationship between peak velocity and amplitude (Fig. 3B) (Zuber et al., 1965). Interestingly, and as we show explicitly in more detail below, we observed the strongest microsaccadic suppression for the white fixation spot condition, which had the smallest, and thus slowest, eye movements. We return to this point later in the text, after describing the subjects’ perceptual performance results in the task.

**Figure 3.**
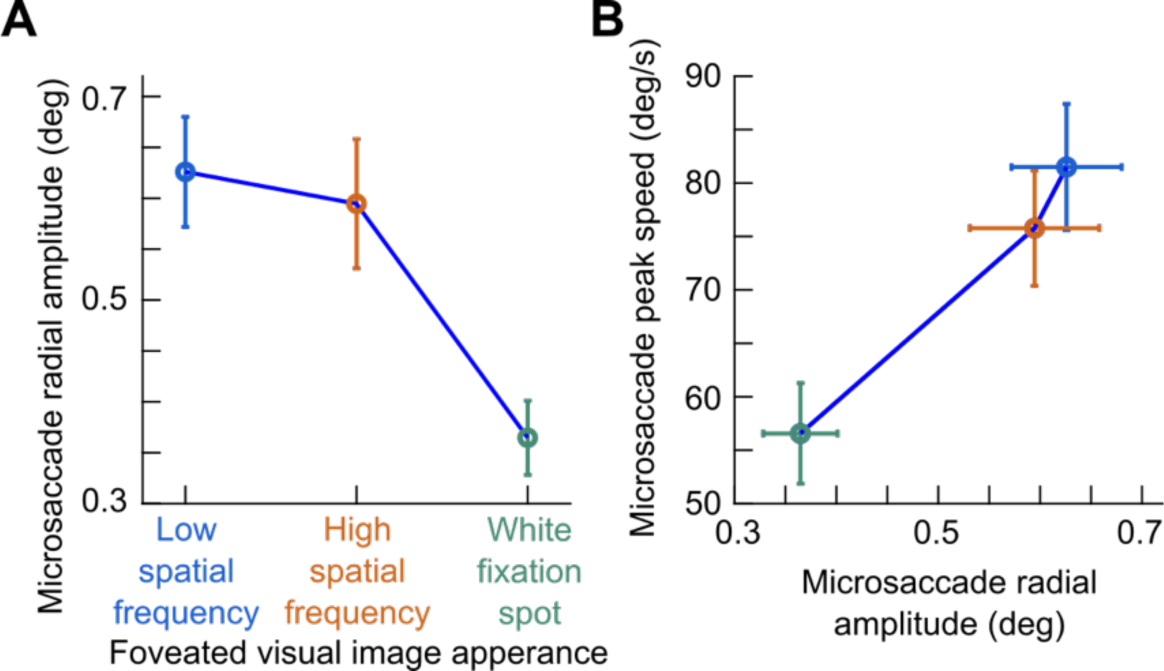
Interaction between foveated visual image appearance and microsaccade amplitudes. (A) For all trials with the nearest microsaccade to probe flash onset being predominantly horizontal, we plotted the radial amplitude of the microsaccade as a function of the foveated image type. Error bars denote SEM across subjects. Microsaccade amplitudes reflected the underlying spatial scale of the viewed foveal image, as expected, but they were always significantly smaller than 1 deg. **(B)** Same as **A** but now plotting the peak velocity of the microsaccades as a function of their amplitude. Error bars denote SEM across subjects. The eye movements followed the expected main sequence relationship between saccade size and saccade peak speed (Zuber et al., 1965). Also see Fig. 8 below for raw microsaccade amplitude distributions across image types.

Thus, given that we have now confirmed the occurrence of small, fixational microsaccades in our experiments, we now had a situation relatively similar to that described in (Idrees et al., 2020): that is, a rapid eye movement (this time, small) was generated across a textured background, and the detection of a brief probe flash away from the saccade end point was investigated. We now turn to describing how the detection of the probe flash varied as a function of both its time relative to microsaccade onset time as well as the underlying foveated visual image appearance. We finish by exploring the potential influences of gaze position and microsaccade amplitude differences on the interpretation of our results.

### Both perceptual detection thresholds and sensitivity are affected in the immediate temporal vicinity of microsaccades

Figure 4 shows psychometric curves characterizing the performance of one example subject (S05) when fixating the low spatial frequency grating. The baseline curve (gray in Fig. 4A, B) was obtained from all trials in which there were no microsaccades occurring within +/- 250 ms from probe flash onset (Methods). For the other shown curves, a predominantly horizontal microsaccade started either within +/- 50 ms from probe flash onset (that is, during the expected microsaccadic suppression time bin; Fig. 4A) or 70 to 150 ms before probe flash onset (that is, during a recovery time bin with the microsaccadic event being sufficiently far away in time; Fig. 4B). As can be seen, the subject’s performance in the task was clearly impaired in the microsaccadic suppression time bin, as evidenced by the lower proportion of correct trials in every flash level that was neither too difficult (floor effect) nor too easy (ceiling effect). For probe flashes longer after a microsaccade (in the recovery time bin), performance recovered and approached that observed in the baseline trials characterized by an absence of nearby microsaccades to the probe flash (Fig. 4B).

**Figure 4.**
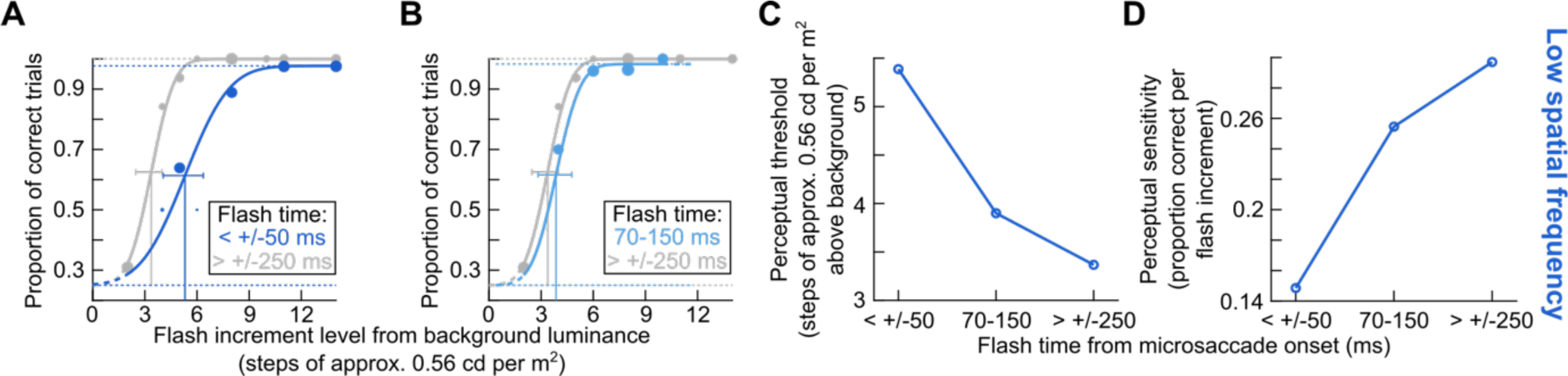
Impairment in both detection threshold and sensitivity (slope of the psychometric curve at threshold) by microsaccades in an example subject. (A) When viewing the low spatial frequency grating (Fig. 1A), peripheral probe flashes (at approximately 9 deg eccentricity) had to be higher in luminance to be successfully detected if they occurred within +/- 50 ms (blue) from microsaccade onset than if they occurred without any nearby microsaccades within +/- 250 ms (gray). This figure shows the results from one example subject (S05). The continuous curves show psychometric curve fits to the shown data points (Methods), and each data point’s size is scaled by the number of observations collected with the shown probe flash luminance increment above the background luminance. The vertical lines show the flash levels resulting in threshold performance (the slight y-value differences at the threshold indications reflect the slightly different asymptotic levels of performance in the two shown conditions) (Schutt et al., 2016). **(B)** Same analyses but with the flashes now occurring 70 to 150 ms after microsaccade onset. Performance recovered to near baseline performance. The gray curve is the same as that in **A**. **(C)** Detection thresholds of this subject as a function of flash time relative to microsaccade onset. There was clear peri-microsaccadic suppression of performance (manifested as a threshold elevation). **(D)** Similar to **C** but for measures of the slope of the psychometric curves near the threshold values (Methods). Psychometric curves were shallower for probe flashes within +/- 50 ms from microsaccade onset.

The impairment of the example subject’s performance in the microsaccadic suppression time bin was manifested in two ways. First, there was an increase in detection threshold. For example, during the microsaccadic suppression time bin (Fig. 4A), the probe flash needed to be almost approximately 3.1 cd/m^2^ brighter than the background to result in a 62.5% correctness rate in task performance (Fig. 4C, leftmost data point). This was a Weber contrast value of 0.14. On the other hand, during baseline, the probe flash needed to be only approximately 1.9 cd/m^2^ brighter than the background luminance (Fig. 4C, rightmost data point); equivalent to a 0.09 Weber contrast. Second, the sensitivity of performance to subtle luminance changes of the probe flash was also impaired. This is evidenced by the shallower slope of the psychometric curve of the subject during the microsaccadic suppression time bin (Fig. 4A) when compared to the recovery and baseline conditions (also quantified in Fig. 4D). In the recovery time bin (Fig. 4B), the slope of the psychometric curve was more similar to that in baseline, suggesting an expected gradual return to baseline sensitivity with time (Fig. 4D). Thus, both the detection threshold and sensitivity (slope of the psychometric curve at the perceptual threshold) of the subject were impaired in association with microsaccades.

Across all subjects, microsaccadic suppression affected both detection thresholds and sensitivity (psychometric curve slopes), and for all foveated visual image appearances that we tested. This is best seen by the analyses of Fig. 5. Here, we plotted in the left column (Fig. 5A, C, E) the detection thresholds of all subjects as a function of probe flash time relative to microsaccade onset time. The different panels denote the different fixated images, and the error bars denote SEM across subjects. In each panel, there was an elevation of perceptual detection thresholds in the microsaccadic suppression time bin, which recovered in other time windows. As for sensitivity (the slope of the psychometric curve at the perceptual threshold flash level), the results are shown in Fig. 5B, D, F. Here, the microsaccadic suppression time bin was associated with generally reduced sensitivity (shallower psychometric curves) relative to the other two analyzed time windows, and this happened for all foveated image types. Statistically, these results were robust. Specifically, within each image type, there was a significant effect of flash time on perceptual thresholds (p=0.0061992, 0.0001872, and 0.00030864 for the low spatial frequency, high spatial frequency, and white fixation spot, respectively; Friedman test with time bin as factor). There were also effects on the slopes of the psychometric curves (p= p=0.024611, 0.00050886, and 0.022274 for the low spatial frequency, high spatial frequency, and white fixation spot; Friedman test with time bin as factor). In post-hoc comparisons, the threshold in the microsaccadic suppression time bin was systematically different from that in the baseline time bin (p=0.0073, 0.0005, 0.0005 for the low spatial frequency, high spatial frequency, and white fixation spot, respectively; Wilcoxon signed-rank test). Similarly, the slopes tended to also be different between the microsaccadic suppression and baseline time bins (p=0.0049, 0.0063, 0.0400 for the low spatial frequency, high spatial frequency, and white fixation spot, respectively; Wilcoxon signed-rank test). Thus, both perceptual detection thresholds and sensitivity (psychometric curve slopes) were affected in the immediate temporal vicinity of microsaccades.

**Figure 5.**
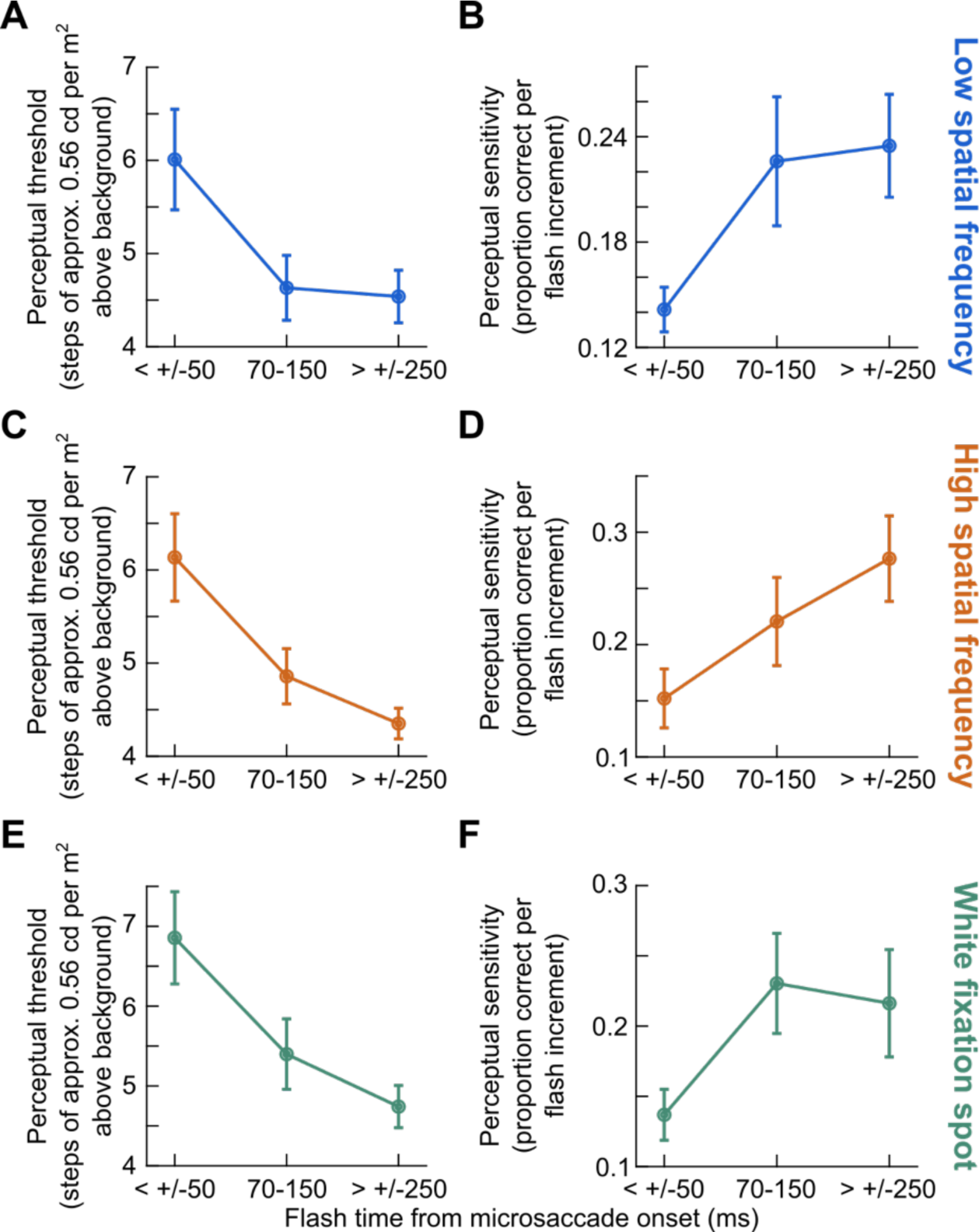
Summary across subjects. (A) Detection thresholds as a function of flash time from microsaccade onset when viewing the low spatial frequency grating. This figure is similar to Fig. 4C, but averaging across all subjects. Error bars denote SEM across subjects. There was a threshold elevation near microsaccade onset, and recovery for larger temporal separation between microsaccades and flash times. **(B)** Same as **A** but for the slopes of the psychometric curves when viewing a low spatial frequency grating. This figure is, thus, similar to Fig. 4D. **(C, D)** Same as **A**, **B**, but for the case of viewing the high spatial frequency grating. Qualitatively similar observations were made. **(E, F)** Same as **A**, **B**, but for the case of viewing the white fixation spot. Again, qualitatively similar observations were made. See Fig. 6 for quantitative comparisons.

### Only perceptual detection thresholds might depend on the foveated image appearance

Despite the qualitatively similar results in Fig. 5 across all three foveated visual image appearances, when we quantitatively compared these results, we found that microsaccadic suppression of peripheral perceptual detection performance was strongest when viewing the small white fixation spot rather than when viewing a low spatial frequency grating, as we had previously observed with large saccades (Idrees et al., 2020). Consider, for example, Fig. 6A, which combines the threshold plots of Fig. 5 together into one single visualization. In the microsaccadic suppression time bin, there was a higher threshold value for the white fixation spot than for the low spatial frequency grating (p=0.01389; Friedman test comparing all three image conditions; and p=0.0161; post-hoc Wilcoxon signed-rank test comparing the white fixation spot to the low spatial frequency grating condition). This effect also continued in the recovery time bin (p=0.0022806; Friedman test; p=0.002; post-hoc Wilcoxon signed-rank text comparing the white fixation spot to the low spatial frequency grating condition), consistent with the higher threshold for the white fixation spot in the microsaccadic suppression time bin. In the baseline time bin, all the detection thresholds were statistically similar to each other (p=0.35255; Friedman test comparing all three image conditions in the baseline time bin). During microsaccadic suppression, the high spatial frequency performance was intermediate between the two, and closer to the low spatial frequency condition. Thus, in terms of detection thresholds, peri-microsaccadic suppression of peripheral perceptual detection performance was strongest for the white fixation spot, as opposed to either a low or high spatial frequency grating.

**Figure 6.**
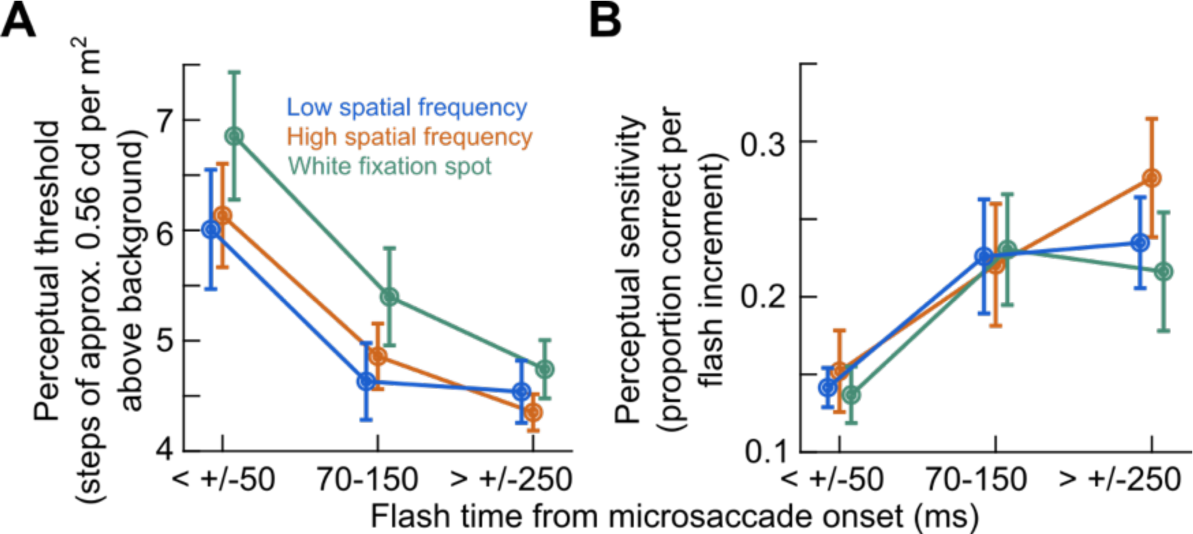
Stronger microsaccadic suppression for the small white fixation spot. (A) We plotted the threshold values of Fig. 5 together in one graph. Subjects exhibited higher detection thresholds for the white fixation spot, suggesting stronger microsaccadic suppression. Moreover, this stronger suppression persisted in the recovery time bin (70-150 ms from microsaccade onset), and it was only in baseline that the thresholds for all three viewed image types were statistically similar. Quantitatively, the thresholds during the microsaccadic suppression time bin were 3.84, 3.44, and 3.36 cd/m^2^, respectively, for the white fixation spot, high spatial frequency, and low spatial frequency grating (equivalent to 0.173, 0.155, and 0.152 Weber contrast, respectively). **(B)** For the slopes of the psychometric curves, there were no differences across viewed image types in any of the time bins. Thus, only detection thresholds showed an image-dependence of microsaccadic suppression in our data.

Interestingly, unlike the thresholds, the slopes of the psychometric curves of the same subjects did not appear to depend on the foveated visual image appearance during the microsaccadic suppression time bin (p=0.59156; Friedman test comparing all three image conditions). This can be seen in Fig. 6B. For all image types, the shallower slope during the microsaccadic suppression time bin was similar in value. This was also the case for the higher slopes seen in the recovery and baseline time bins. Thus, it was only the detection thresholds, and not the slopes of the psychometric curves, that showed an image-dependence of microsaccadic suppression of peripheral perceptual detection performance. It would be interesting in future studies to investigate why this was the case.

We next considered whether the results of Fig. 6A could be explained by factors other than microsaccadic suppression per se. Specifically, since there was no specific marker to fixate on in the grating images, it could be the case that the subjects were biased in where they directed their gaze during the trials with grating images. For example, if these subjects systematically fixated their gaze slightly upward relative to the grating center, then this could have rendered one flash location (the upper one in this example) significantly closer to the gaze center than in the case of the small white fixation spot, with much more focused gaze direction. This would have made one flash location easier to detect than with the white fixation spot. However, this logic fails since a bias in gaze position with the gratings towards one probe flash location would render the three other flashes actually farther away from the gaze center, and therefore harder to detect. If anything, this should have made the overall task harder with the grating images than with the white fixation spot. This was clearly not the case in our data (subjects performed worse during the microsaccadic suppression time bin with the white fixation spot). We also have three additional reasons to rule out a potential influence of gaze position (and thus probe flash visibility) on the results of Fig. 6A.

First, we explicitly measured gaze position at the time of probe flash presentation across all trials and image types (Methods). Figure 7 shows these measurements for each subject individually, with the blue dots showing trials with the low spatial frequency grating and green dots showing trials with the white fixation spot. We did not plot the high spatial frequency grating data in order to reduce clutter in the figure, but these data were virtually identical to those of the low spatial frequency grating data (consistent with Fig. 3). The insets show the mean and standard deviations of the shown raw data points. As can be seen, while it was certainly true that gaze position was more dispersed with the grating images, as expected, the subjects correctly followed our instructions to maintain their gaze near the center of the image (average gaze position was similar whether the subjects were viewing a grating or a small white fixation). Quantitatively, deviations in mean gaze position between the white fixation spot and the grating cases were always smaller than approximately 0.5 deg, and often significantly smaller. Given that our peripheral probes were at 9.1 deg, this small difference in average gaze position was not expected to influence detectability. In fact, even at 5 deg, we found in an earlier study that such gaze position deviations of the same magnitude as those observed here did not alter peripheral detection performance (J. Bellet et al., 2017).

**Figure 7.**
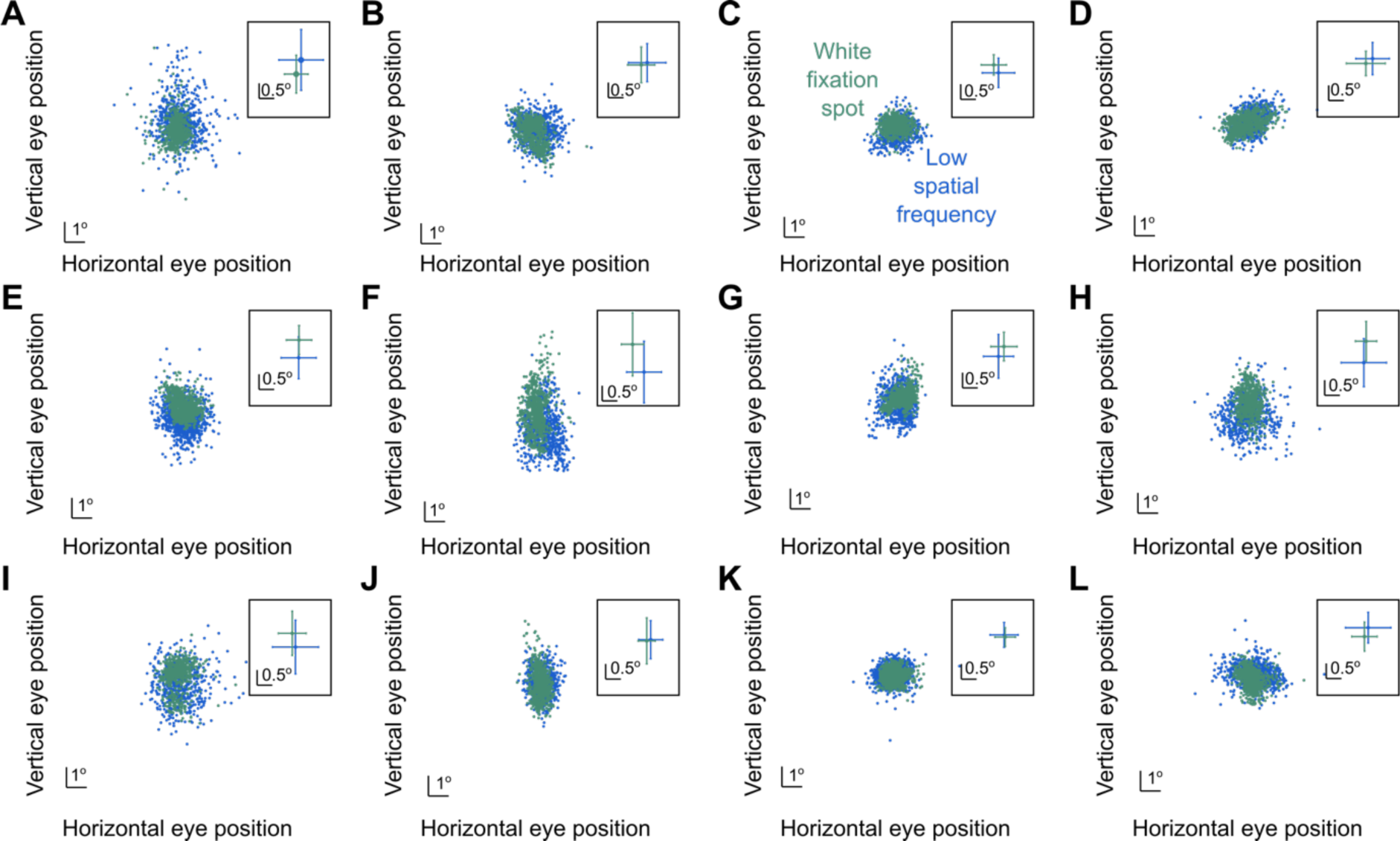
Lack of large systematic gaze position biases with grating images. For each subject (each panel), we measured eye position at the time of probe flash presentation (Methods). In each panel, each dot represents a single trial, and the different colors indicate which image was being viewed by the subject (the color legend in **C** applies to all panels). The inset in each panel shows the mean and standard deviation values of the corresponding raw data plots of the panel, and **E** contains the eye positions of the same example subject whose psychometric curves were shown in Fig. 4 above (S05). As can be seen, all subjects fixed their gaze at very similar positions between the two image types. There was clearly larger dispersion of fixation position with the grating images (due to a lack of a specific punctate marker), but this dispersion was largely symmetric in all directions. Thus, all four peripheral flashes were, on average, at a similar retinotopic eccentricity when they appeared, ruling out a simple retinotopic visibility as the primary explanation of the results of Fig. 6A.

Second, in the baseline time bin in Fig. 6A, perceptual performance was similar for all image types. Thus, if gaze position was indeed systematically biased for the gratings relative to the white fixation spot, then we should have also seen a difference in performance during the baseline time bin. This was not the case.

Third, the stronger microsaccadic suppression of peripheral detection performance for the white fixation spot (as opposed to the grating images) was specific for threshold values but not for the slopes of the psychometric curves (Fig. 6B). If gaze position altered the visibility of the peripheral targets between the different image types, then we might have expected similar changes in both thresholds and sensitivity across image types. Therefore, all of these observations, coupled with the results of Fig. 7, suggest that systematic gaze position differences across image conditions likely do not fully explain the results of Fig. 6.

This leaves a final question of microsaccade size itself; that is, it could be possible that microsaccade size could influence the results of Fig. 6A. In Fig. 3, we did indeed observe that microsaccades were smaller and slower, on average, with the white fixation spot when compared to the grating images. However, slower movements should cause milder image transients and blurs than faster movements, which should, in principle, be associated with milder saccadic suppression. This is opposite to what we observed experimentally, with stronger microsaccadic suppression for the white fixation spot condition. Moreover, there seems to be a dissociation between saccade speed and saccadic suppression strength in general (Gremmler & Lappe, 2017). And, an early study with large saccades actually documented larger saccadic suppression effects with larger (and faster) saccades (Mitrani, Yakimoff, & Mateeff, 1970), again opposite of what we observed. Nonetheless, we investigated our microsaccade amplitude distributions more closely. Despite the differences in average microsaccade sizes that we observed in Fig. 3, there was a large overlap in the raw distributions of microsaccade sizes, as can be seen from Fig. 8 (with most microsaccades being smaller than 1 deg in all conditions). Thus, it does not seem likely that the results of Fig. 6A could be fully accounted for by microsaccade size, and we confirmed this (in a control analysis) by excluding all grating image trials containing microsaccades larger than 1 deg in amplitude. This maximized the overlap in microsaccade sizes across all image conditions, and we observed similar trends as those shown in Fig. 6. It would be interesting in future experiments to relate, within a single image, microsaccadic suppression strength to the sizes of the microsaccades that are generated, in order to explicitly document whether it was the smaller microsaccades in the white fixation spot condition that fully accounted for our results or not.

**Figure 8.**
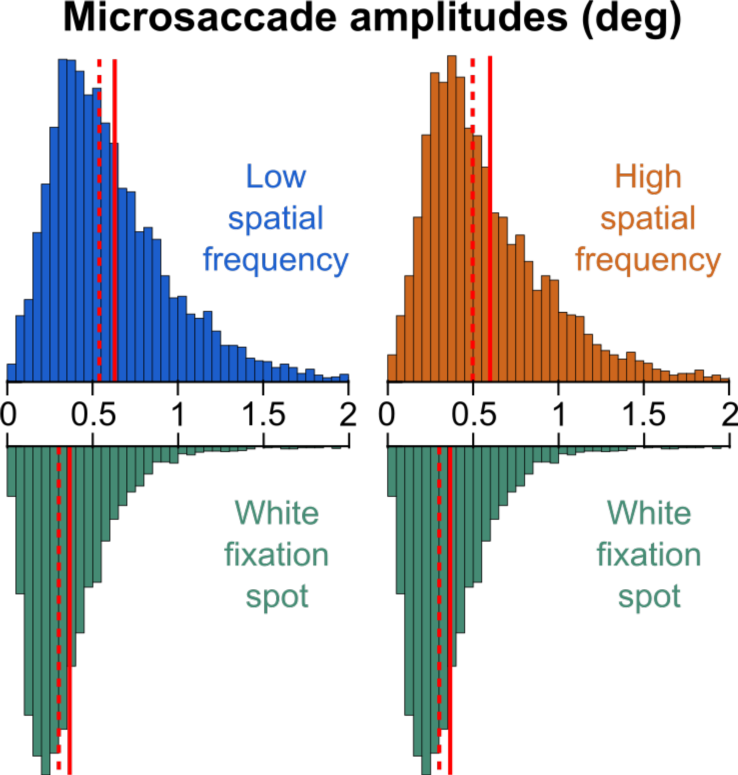
Microsaccade amplitude distributions. In this figure, we plotted the raw microsaccade amplitude distributions underlying the summary statistics in Fig. 3. We placed the distribution for the white fixation spot under each of the low or high spatial frequency gratings for easier comparison. In all cases, most microsaccades were smaller than 1 deg in amplitude. Thus, there was large overlap across conditions. In each distribution, the dashed vertical line indicates the median, and the solid vertical line indicates the mean.

Therefore, our analyses, combined, suggest that microsaccadic suppression in our experiments did occur for all tested visual image types at the fovea, and that the suppression was stronger when the viewed foveal visual image was a small white fixation spot as opposed to either a low or high spatial frequency texture.

## Discussion

In this study, we investigated the dependence of peri-microsaccadic suppression of peripheral perceptual detection performance on the visual appearance of the images across which microsaccades were generated. This was a microsaccadic correlate of studies in which saccadic suppression was researched with larger saccades being made across textured backgrounds (Idrees et al., 2020). Unlike a potential expectation from these studies, we did not find stronger microsaccadic suppression for the low spatial frequency grating. Rather, the strongest suppression occurred when microsaccades were made across a small, foveal fixation spot (Fig. 6A). Moreover, only detection threshold elevations depended on the foveal visual image appearance, but not the slope reductions of the psychometric curves.

Our effect on threshold elevations, being slightly larger for the small white fixation spot than for gratings (Fig. 6A), was quite different from what we observed earlier with much larger saccades (Idrees et al., 2020), in which there was stronger saccadic suppression for low than high spatial frequency background textures. This difference in results might be a manifestation of the significantly slower and smaller eye movements studied here when compared to our earlier experiments. For example, with the small eye movements, shifting gaze on the low spatial frequency grating might be more similar to shifting gaze over a blank, given how gradually the grating luminance changes with the diminutive angular displacements associated with microsaccades. Similarly, with the high spatial frequency grating, the image displacement caused by the average microsaccade size that we observed (approximately two thirds of a degree; Fig. 3) might have shifted the image by a whole number multiple of luminance cycles (e.g. 3 cycles), just averaging the luminance modulations across the cycles out by the time the eye movement was finished. On the other hand, with the broad band fixation spot, shifting gaze over it would be expected to influence multiple spatial frequency visual processing channels, and might cause stronger saccadic suppression. This interpretation is consistent with the idea that saccades of different sizes have different spatio-temporal profiles of retinal image modulations when they occur (Mostofi et al., 2020). Thus, the interactions between background image spatial frequency content and the strength of saccadic suppression (Idrees et al., 2020) should not always be identical for different saccade sizes; rather, these interactions might reflect the specific sensory consequences of the particular saccades being generated.

Given that microsaccades to a small spot can overshoot it slightly (Tian et al., 2016; Willeke et al., 2022; Willeke et al., 2019), making a microsaccade in our white fixation spot condition was additionally equivalent to crossing a luminace edge (e.g. the preferred retinal locus went from a gray background to being over a white image patch and then to being over a gray background again by the end of a given microaccade). This is a similar situation to our recent observation that when large saccades crossed a luminance bar, we observed stronger saccadic suppression than when the saccades were made across a blank (Baumann et al., 2021). Thus, a second potential explanation of the results of Fig. 6A is that the microsaccades with the white fixation spot were crossing a luminance discontinuity. This would still be consistent with a visual component to microsaccadic suppression, as with larger saccades. However, in the current study, the receptive fields experiencing the peripheral probe flashes (especially if they were small in early visual areas like retina, lateral geniculate nucleus, and primary visual cortex) likely never crossed luminance bars when the microsaccades happened with the white fixation spot; these receptive fields presumably always experienced a gray background since the probe flashes were at a peripheral eccentricity. Thus, it is not clear whether in our earlier study (Baumann et al., 2021), it was the foveal crossing or the crossing of the receptive fields seeing the probes of a luminance bar that ultimately caused the stronger saccadic suppression. It would be interesting in the future to investigate this issue further.

Indeed, all descriptions above include an implicit assumption that the visual conditions in the fovea in our current experiments could influence peripheral performance even though the peripheral probe flashes themselves occurred over a completely gray background. However, this is not the first time that probes over a gray background were shown to be affected by visual conditions far from them. For example, in some experiments in (Idrees et al., 2020), we had probe flashes over a gray background (with the same retinal locus being stimulated by gray both before and after saccades), and only the far surround having different textures. We still obtained altered saccadic suppression strengths with the differing far surrounds. Thus, it is still possible that our results in Fig. 6A could be affected by the foveal image even though the probes were peripheral. Indeed, the thresholds that we observed in the current study during the microsaccadic suppression time bin (e.g. 0.173 Weber contrast for the white fixation spot) were slightly higher than those observed in (Baumann et al., 2021) with saccades made across a completely blank background in the same experimental setup (0.13 Weber contrast). Thus, it could be the case that the foveal visual conditions in the current study could still influence peripheral performance over a gray background.

This leads to the intriguing question of how and why peripheral sensitivity can be impaired so much when tiny microsaccades occur. In the above example, our perceptual thresholds with the white fixation spot were slightly higher than those we observed earlier with much larger saccades over a gray background (Baumann et al., 2021). This also happens at the neuronal level, with even very eccentric receptive fields (preferring >20 deg of eccentricity) experiencing massive suppression of visual neural sensitivity to probe onsets whenever tiny microsaccades occur (Chen & Hafed, 2017; Hafed, Chen, & Tian, 2015; Hafed & Krauzlis, 2010). Of course, one aspect of this could still be visual. For example, at least in the superior colliculus, a structure relevant for saccadic suppression (Berman et al., 2017; Lee et al., 2007; Phongphanphanee et al., 2011), eccentric receptive fields can be very large. Thus, moving them by microsaccadic amounts can still cause luminance transients associated with the display edge moving on the retina relative to the dark surroundings of the display region. That is, a single peripheral receptive field (e.g. in the superior colliculus) can still experience both the display and the dark background within it, and thus be exposed to a visual edge movement whenever tiny microsaccades occur.

There could also be other non-visual components for such a disparate difference between saccade size and the peripheral eccentricity that experiences suppression. For example, if microsaccade-related motor bursts in the superior colliculus were to vary as a function of the image appearance, as is the case with larger saccades (Baumann et al., 2023), then a dependence of microsaccadic suppression on foveal visual image appearance could emerge peripherally through extra-retinal mechanisms (with concepts like corollary discharge). It seems likely that microsaccade-related superior colliculus motor bursts would exhibit image-dependence given the current evidence in the literature so far. For example, these motor bursts can disappear completely for microsaccades made towards a blank (Willeke et al., 2019), just like with larger saccades (Baumann et al., 2023; Edelman & Goldberg, 2001; Mohler & Wurtz, 1976; Zhang et al., 2022). However, an explicit experiment probing collicular microsaccade-related motor discharge with different underlying foveal textures is warranted.

Finally, it would be interesting in follow-up experiments to present our probe flashes more centrally, such that they still appear on the low or high spatial frequency gratings themselves, and not over the gray background. In that case, we can expect higher overall detection thresholds than with a blank background (Baumann et al., 2021), but it remains to be seen whether microsaccadic suppression of performance would now be stronger for the low spatial frequency grating than for the high spatial frequency grating. In such experiments, one can even parametrically change grating size relative to probe flash location, in order to find the extent of overlap between background images and probe flashes that is needed to result in an image-dependence of microsaccadic/saccadic suppression. This can turn allow predicting the effective sizes of receptive fields that would be most relevant for the visual component of saccadic and microsaccadic suppression (e.g. in retina, lateral geniculate nucleus, primary visual cortex, superior colliculus, or elsewhere).

Overall, we believe that our results demonstrate the relevance of studying saccadic suppression using different eye movement sizes and directions, and also the relevance of considering both visual and motor components of this highly ubiquitous and robust perceptual phenomenon.

## Acknowledgements

We were funded by the Deutsche Forschungsgemeinschaft (DFG): SFB 1233, Robust Vision: Inference Principles and Neural Mechanisms, project 11, project number: 276693517. We also thank Matthias P. Baumann for helpful comments on the manuscript.

## Notes

### Competing Interest Statement

The authors have declared no competing interest.

